# The omicron (B.1.1.529) SARS-CoV-2 variant of concern does not readily infect Syrian hamsters

**DOI:** 10.1101/2021.12.24.474086

**Authors:** Rana Abdelnabi, Caroline S. Foo, Xin Zhang, Viktor Lemmens, Piet Maes, Bram Slechten, Joren Raymenants, Emmanuel André, Birgit Weynand, Kai Dallemier, Johan Neyts

## Abstract

The emergence of SARS-CoV-2 variants of concern (VoCs) has exacerbated the COVID-19 pandemic. End of November 2021, a new SARS-CoV-2 variant namely the omicron (B.1.1.529) emerged. Since this omicron variant is heavily mutated in the spike protein, WHO classified this variant as the 5th variant of concern (VoC). We previously demonstrated that the other SARS-CoV-2 VoCs replicate efficiently in Syrian hamsters, alike also the ancestral strains. We here wanted to explore the infectivity of the omicron variant in comparison to the ancestral D614G strain. Strikingly, in hamsters that had been infected with the omicron variant, a 3 log_10_ lower viral RNA load was detected in the lungs as compared to animals infected with D614G and no infectious virus was detectable in this organ. Moreover, histopathological examination of the lungs from omicron-infecetd hamsters revealed no signs of peri-bronchial inflammation or bronchopneumonia. Further experiments are needed to determine whether the omicron VoC replicates possibly more efficiently in the upper respiratory tract of hamsters than in their lungs.

## Main text

Variants of SARS-CoV-2 are still emerging in different parts of the world, posing a new public health threat. Even in highly endemic regions, some of these variants have replaced the formerly dominant strains and resulted in new waves of infections and new spikes in mortality *(1)*. On 24 November 2021, South Africa officially reported the emergence of B.1.1.529 (omicron) variant to WHO. Two days later, the omicron variant has been classified by WHO as the 5th variant of concern (VoC) following the alpha, beta, gamma and delta VoCs *(2)*. Among these VoC, the omicron variant carries the highest number of spike protein mutations (>30 mutations) *(3)*. Some of the spike mutations carried by the omicron variant have been reported in other VoCs to be associated with immune escape and reduced susceptibility to monoclonal antibodies *(3)*. In addition, the omicron variant carries some spike mutations that could be involved in increased transmissibility, which is also supported by the rapid replacement of delta variant by omicron as the dominant variant in South Africa *(3)*. Currently, there is not enough clinical data to indicate whether the Omicron variant can cause more severe disease. We previously showed that alpha, beta, gamma and delta VoCs are replicating efficiently in the lungs of Syrian hamsters in similar extent to the ancestral strains (i.e. Wuhan and D614G strains) *(4–6)*. We here compare the infectivity of the omicron variant versus the ancestral D614G strain in our Syrian hamster model. The ancestral strain used in this study is strain Germany/BavPat1/2020 (also referred to as BavPat-1, EPI_ISL_406862; 2020-01-28)*(7)*. This strain carries a spike D614G substitution found in early European variants and linked to more efficient transmission *(8)*. The omicron (B.1.1.529) variant was isolated from a nasopharyngeal swab taken from a traveler returning to Belgium in end of November 2021 (hCoV-19/Belgium/rega-20174/2021, EPI_ISL_6794907).

In Brief, 6-8 weeks old female Syrian hamsters were intranasally infected with 50 µL containing approximately 10^3^ TCID_50_ of either the ancestral strain (BavPat(D614G)) or the omicron VoC (B.1.1.529) SARS-CoV-2 (Fig. 1a) as described previously *(9, 10)*. At day four post-infection (4 dpi), animals were euthanized for sampling of the lungs and further analysis by i.p. injection of 500 μL Dolethal (200 mg/mL sodium pentobarbital) *(4)*. Housing conditions and experimental procedures were approved by the ethics committee of animal experimentation of KU Leuven (license P065-2020).

**Fig. 1.**
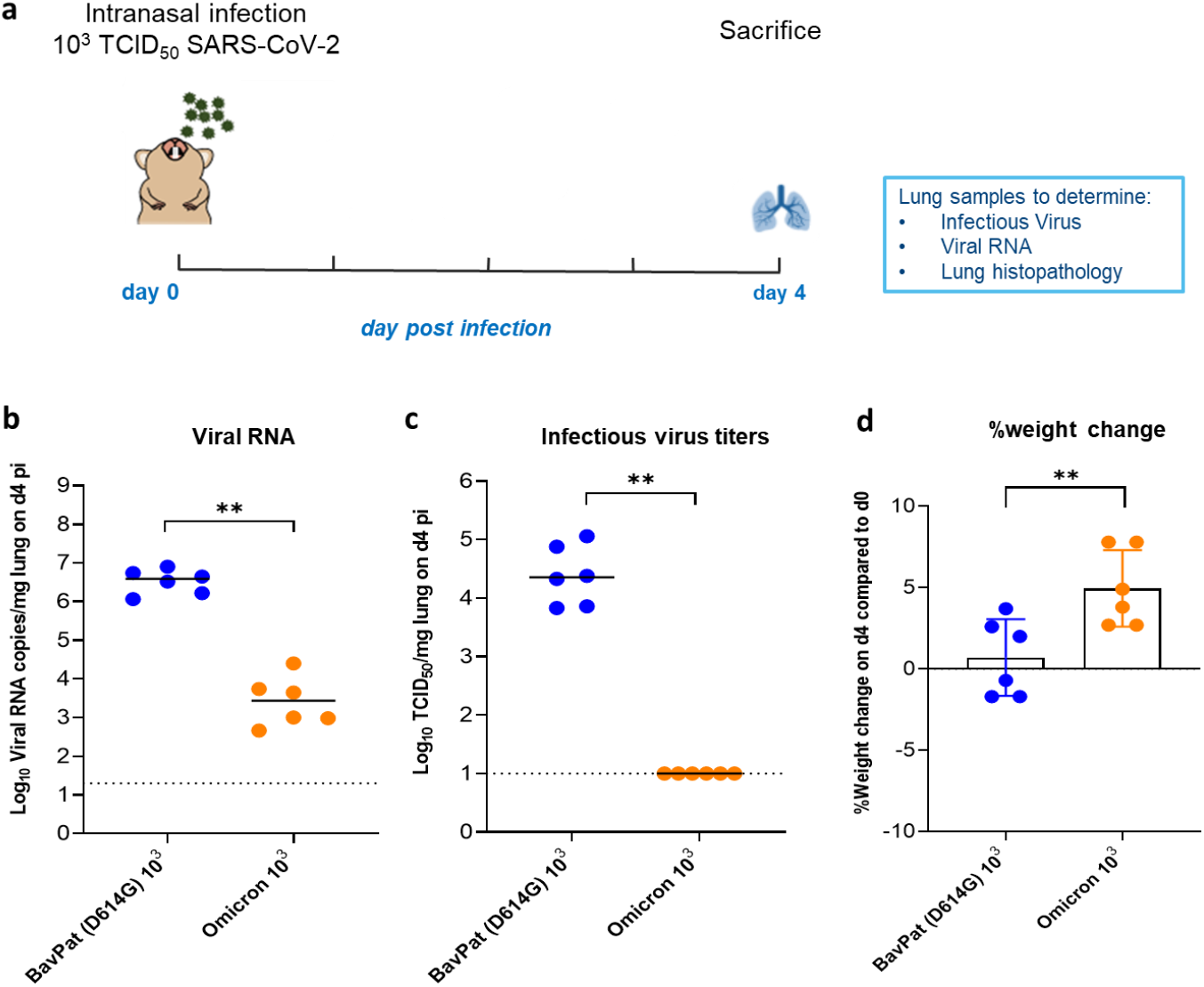
Characterization of the *in vivo* replication of the omicron SARS-CoV-2 variant versus the ancestral D614G strain. (a) Set-up of the Syrian hamster infection study. (b) Viral RNA levels in the lungs of hamsters infected with 10^3^ TCID_50_ of BavPat (D614G) strain (n=6) or the omicron (B.1.1.529) SARS-CoV-2 variant (n=6) on day 4 post-infection (pi) are expressed as log_10_ SARS-CoV-2 RNA copies per mg lung tissue. Individual data and median values are presented. (c) Infectious viral loads in the lungs of hamsters infected with the D614G strain or the omicron variant at day 4 pi are expressed as log_10_ TCID_50_ per mg lung tissue. Individual data and median values are presented. (d) Weight change at day 4 pi in percentage, normalized to the body weight at the time of infection. Bars represent means ± SD. Data were analyzed with the Mann–Whitney U test, **P < 0.01. All data are from a single experiment.

A median Viral RNA load of 4×10^6^ RNA copies/mg of lung tissue was detected at 4 dpi in the lungs from the animals infected with the D614G strain (**Fig. 1b**). On the other hand, ∼3 log_10_ lower viral RNA loads were detected in the lungs of animals infected with the omicron variant (a median vira RNA load of 3×10^3^ RNA copies/mg lung tissue, p=0.0022, Mann-Whitney Test), **Fig. 1b**. Infectious virus titers in the lungs of D614G strain-infected animals were around 2×10^4^ TCID_50_/mg of lung tissue (**Fig. 1c**). Strikingly, no infectious virus titers were detected at all in the lungs of all the animals infected with the omicron variant (**Fig. 1c**, P=0022 compared to the D614G strain-infected group, Mann-Whitney Test). This is also different from the other four VoCs which proved to replicate efficiently and consistently to high viral loads in Syrian hamster lungs by day 4 post-infection *(4–6)*. On the day of sacrifice, animals infected with the omicron variant showed more increase in body weight (average body weight change from d0 of 3.8%) than the D614G strain-infected animals (average body weight change from d0 of 0.65%), p=.0087, Mann-Whitney Test (**Fig. 1d**).

Hematoxylin/eosin (H&E)-stained images of lungs of hamsters infected with the D614G strain showed significant pathological signs including peri-bronchial inflammation, bronchopneumonia in the surrounding alveoli and perivascular inflammation with peri-vascular oedema (Fig. 2a). The median cumulative histopathological lung score of the D614G-infected hamsters was 7.5 (Fig. 2b), which is similar to what we previously reported for this strain *(4)*. Unlike the D614G strain-infected group, no inflammation or disease signs were observed in the lungs of the omicron-infected animals on day 4 pi (Fig. 2a). The median cumulative histopathological lung scores of the omicron-infected animals was close to the baseline score in untreated, non-infected hamsters (median score of 1.75, Fig. 2b, P=0022 compared to the D614G group, Mann-Whitney Test).

**Fig. 2.**
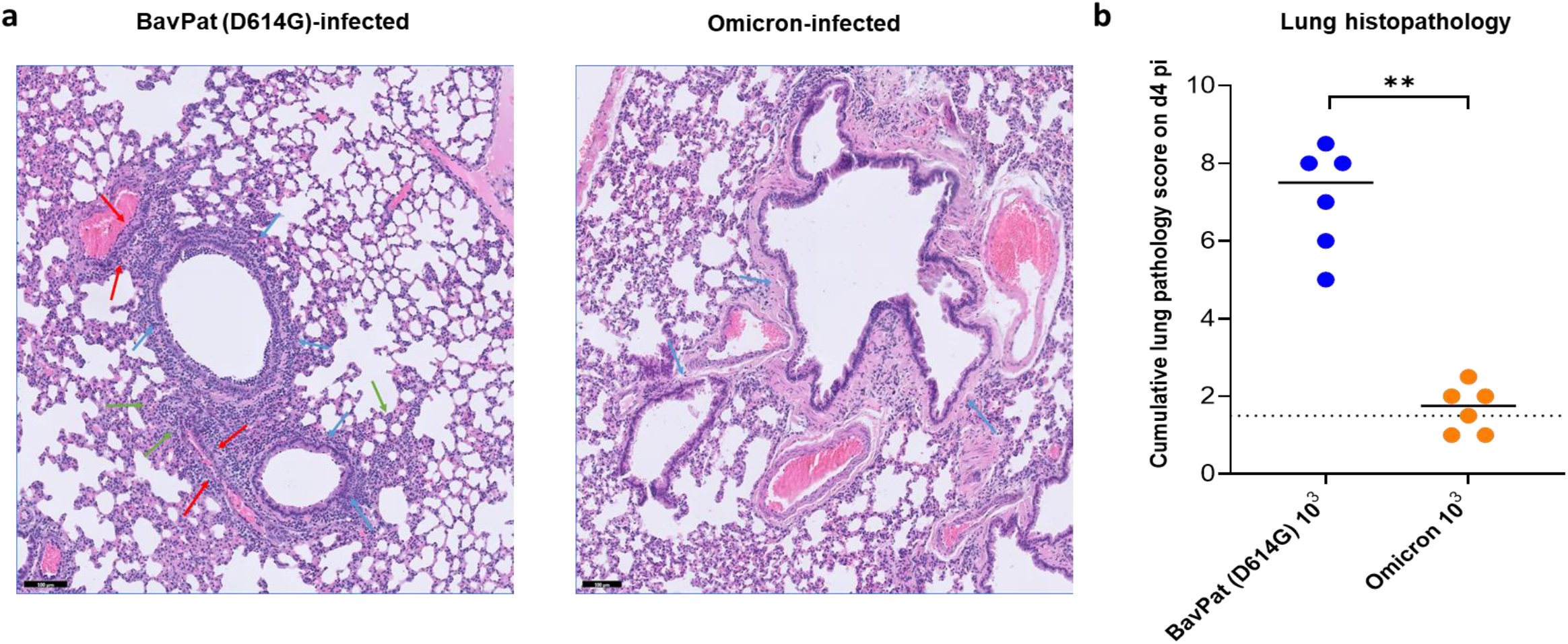
Histopathology of lungs of Syrian hamsters infected with either the D614G strain or the omicron SARS-CoV-2 variant. (a) Representative H&E images of lungs of hamsters infected with 10^3^ TCID_50_ of BavPat (D614G) strain (n=6) or the omicron (B.1.1.529) SARS-CoV-2 variant at day 4 post-infection (pi). The lungs of hamsters infected with the ancestral D614G strain (left picture) show significant bronchopneumonia (green arrows), perivascular inflammation with peri-vascular oedema (red arrows) and peri-bronchial inflammation (blue arrows), whereas the lungs of the omicron-infected hamsters (Right picture) appear normal with no peri-bronchial inflammation (blue arrows) or bronchopneumonia. Scale bars, 100 μm. (b) Cumulative severity score from H&E stained slides of lungs from hamsters infected with the D614G strain or the omicron variant at day 4 pi. Individual data and median values are presented and the dotted line represents the median score of untreated non-infected hamsters. Data were analyzed with the Mann–Whitney U test, **P < 0.01. All data are from a single experiment.

Taken together, these results clearly demonstrate that the omicron variant is not able to replicate efficiently in the lower respiratory tract of Syrian hamsters compared to the anscetral D614G strain and other variants of concerns when animals were euthanized at day 4 post-infection.

One possible explanation may be that the heavily mutated spike of the omicron variant has now better adaptation to the human ACEII and hence making the attachment of this variant to the hamster ACEII to be less efficient. Another possibility is that the omicron variant tropism could be shifted to the upper respiratory tract resulting in limited lung infectivity. This could be in line with the recentely released ex-vivo models data in which the omicron variant is 70 times more efficient in replicating in human bronchus tissues than the delta variant whereas it is less efficiently replicating in human lung tissues *(11)*. Therefore, further experiments are required to assess the viral loads in lung and other tissues from the upper respiratory tract of omicron-infected hamster at different time points post-infection to explain the limited lung infectivity observed in this study.

## Acknowledgements

We thank Carolien de Keyzer, Lindsey Bervoets, Thibault Francken, Niels Cremers, Jasper Rymenants and Stijn Hendricks for excellent technical assistance with animal experimentation and molecular sample analysis. We thank Bram Van Holm for is technical assistance. We thank Lize Cuypers, Guy Baele, Simon Dellicour and the Belgian Genomic Surveillance program which contributed to the early detection of Omicron in Belgium. We also thanks Bo Corbeels and Kathleen Van den Eynde for histopathology samples processing and staining.

